# In-silico pharmacophore and Molecular Docking based drug discovery against Marburg Virus’s Viral Protein 35; A potent of MAVD

**DOI:** 10.1101/2021.07.01.450693

**Authors:** Sameer Quazi, Shreelaxmi Gavas, Javed Ahmad Malik, Komal Singh Suman, Zeshan Haider

**Affiliations:** GenLab BioSolutions Private Limited, Bangalore, Karnataka, India; Department of Zoology, Guru Ghasidas Vishwavidyalaya, Bilaspur, Chattisgarh, India; Center of Agricultural Biochemistry and Biotechnology, University of Agriculture, Faisalabad, Pakistan

**Keywords:** Marburg virus disease (MVD), Drug Repurposing, Molecular Docking, ZINC database, virtual screening, MARV VP35

## Abstract

Marburg virus is a member of filoviridae and spreads severe Marburg hemorrhagic illness in humans and animals. Nowadays, there is no vaccine available that can completely stop the replication of Marburg replication. Therefore, this study is designed to repurpose the effective therapeutic antiviral drug by using a computational approach against exploring the mechanism of Marburg virus Viral protein 35. We have retrieved about 40570 drug-like small compounds from the ZINC database using the “ZINC Pharmer” online tool. Molecular docking of the ligands from the ready-to-dock database has been carried out using MOE. The five drugs have been identified to bind with VP35 possibly. A study was also performed to evaluate the drug-like characteristics of the substances for absorption, distribution, metabolism, and excretion (ADME). The findings clearly showed that ligands are interacting with the MARV VP35 protein. Interestingly, Lipinski’s rule of five was observed by all ligands. These findings provide the foundation for reconstituting and utilizing molecules as a possible therapy for Marburg Virus Disease (MVD).

## INTRODUCTION

Marburg Virus (MARV) is a unique member of the Filoviridae family of unsegmented negative RNA. MARV infections have a severe hemorrhagic fever, leading to up to 90% mortality in certain instances (1). Anti-filoviral treatment is essential since the filoviruses are aggressive. However, these viruses only encode one protein, an ample enzyme protein (L). An essential approach to create antivirals is thus to aim at non-enzymatic viral proteins utilizing tiny molecules. However, it is challenging to target viral proteins and frequently needs a profound structural and functional understanding of the protein structure and interactions.

The ability of MARV to effectively undermine the host’s innate immune response and subsequent adaptive immune responses has contributed in part to the high mortality rates (2). Among the methods used are the production of interferon (IFN) -α / β and the antiviral signal generated by IFN-α / β (2,3). In particular, the elimination of IFN-α / β is essential for the pathogenicity of MARV (4). The use of various methods of immunosuppression of filoviruses emphasizes the importance of the targeted viral components.

Filoviral VP35 is an intriguing therapeutic target since protein performs many essential functions for virus reproduction, and its structure has been described with remarkable accuracy (5,6). It’s a multifunctional protein that functions as a cofactor in the viral polymerase complex. It may suppress the host’s immunological response, including the generation of interferon (IFN) activated by type I receptors in the retinoic acid-inducible gene (RLR) (RIG-I). The N-terminus of VP35 contains a spiral domain, while the C-terminus has an IFN inhibitor domain (7,8). Recombinant viruses with VP35 mutations are much weaker in guinea pigs and protect against infection with MARV (9). The virus’s development was also reduced by RNA interference against VP35. All of these findings point to VP35’s therapeutic potential (10).

VP35 is a crucial target for developing antiviral medicines since it plays such an essential role in MARV transcription. In the development of drugs, computerized techniques and virtual screening perform a substantial role. Virtual high-throughput screening (vHTS) is an essential and appropriately recognized factor of drug development (11). To block the catalytic centre, MARV VP35, a new cheminformatics protocol, i.e. structure-based ligand screening, molecular docking, and simulation, may be utilized to discover the new drugs compounds in computational biology. Therefore, Using these computational biology methods, the current study aims to find drugs with solid interactions, considerable binding energies, and significant inhibitory efficacy against MARV VP35. These drug-like chemical compounds have a higher probability of inhibiting MARV replication, leading to a MARV cure.

## Methodology

### VP35 structure prediction

The VP35 protein sequence was obtained from the NCBI database. A website called Expasy-Protparam was used to examine the main structure. The homology-based MODELLER v9.25 programme was used to estimate the three-dimensional structure of VP35. Based on the DOPE score, the best model was chosen. Utilizing the electronically tools PROCHECK (https://servicesn.mbi.ucla.edu/PROCHECK/), the quality of the 3D structure of VP35 was assessed and verified. The PROCHECK programme emphasizes protein stereochemistry (12–14).

### Pharmacophoric based virtual screening from ZINC database

The pharmacophoric steric and digital characteristics need to be modified to provide optimum molecular interactions and restrict biological activity with the organic target. The ZINC-pharmer software simulated a study of the active substances in the online ZINC database (15). ZINC Pharmer has taken use of MARV VP35, three-dimensional structure, and the diffusion ratio of “SDF” inhibitor FGI-103 to build pharmacophore-ligand-receptor characteristics, and the promotion of the use of comparative testing requires this kind of screening (16).). Lipinski’s law was utilized to build a drug-like property. Under those requirements, drug-like characteristics should have a log P of five, a molecular weight of 500 da, and five H-binding (HBA) and H-binding (HBD) acceptors. The general rule explains the molecular characteristics of medication in the human body required for pharmacokinetics (17).

The findings of the primary link were chosen and stored for future relationship investigations in a separate file. With the Molecular Operating Environment (MOE) Program, the three-dimensional test database was built.

### Molecular docking

To further explore these drug-related chemicals, all the identified compounds were connected to MARV VP35. MOE methods were used as a standard for removing ligand, hydrogen insertion, and energy reductions of the VP35 protein (18). The search method for the MOE package location was utilized to find the VP35 catalytic centre. The 2D interaction analysis method was utilized to detect and assess ligand-VP35 interaction. Many factors were used to establish molecular interaction between the ligands and VP35 (reactivation function one and reactivation function two: London dG, position: triangular comparator, hold: 2 and refinement: pressure field). The new medications were selected based on the FGI-103 inhibitor score and root means square deviation (RMSD). The S value specifies how much the ligand’s affinity for the recipient is measured by utilizing the usual MOE scoring method. The RMSD technique is used to compare the conformation of the confirmation to the conformation relative to the reference. Compounds having the lowest S and RMSD values were selected for future study (19). The binding energy of these chemicals was estimated using MOE software for future study. The main characteristics of the practical impact of the small molecule on the target protein are the hydrophobic and polar interactions between the ligand and the target protein receptor and its binding affinity to the ligand. Its range of 5 to 15 kcal/mol indicates that the ligand and receptor are strongly interacting with one another (20). The same affinity or more binding chemical according to the classification of reference inhibitors to evaluate the absorption, release, metabolism, excretion, and toxicity profiles further. Small medicinal compounds were also assessed utilizing the basic and biological properties of the pkCSM server.

### Validation and ADMET analysis

The absorption, digestion, metabolism, excretion, and toxicity (ADMET) characteristics of chosen target drugs were evaluated using the pkCSM server (21).

## Result and discussion

In the pursuit of a MARV vaccine, many experimental trials have been conducted. Currently, no medication is being developed to control this infection (11) effectively. And as a consequence, finding a low-expense antiviral medication that suppresses MARV is crucial. One method to ensure that novel medication-like compounds effectively against viral infections is to use a step-by-step approach to drug development. Drug testing in the traditional sense is unreliable and time-consuming (22). In vitro treatment for MARV is presently done using FGI-103, a broad-spectrum antiviral medication. In a computer simulation, the FGI-103 inhibitor of choice also reduced VP35’s biological activity by inhibiting its online active catalyst website (23). It is the most recent development in the search for medicines that may inhibit VP35’s proteolytic activity in the MARV catalytic region.

### VP35 structure prediction

The MARV VP35 virus has 329 amino acids in its viral protein. According to the physicochemical study, the MW of VP35 was 36144.74 Da, with a GRAVY and IN (instability index) of -0.186 and 41.59, respectively. The three-dimensional structure of VP35 is also anticipated using homology-based programme modeller 9.25. The VP35 and the template “ c4gh9A” have a total sequence similarity of approximately 95 percent. The most acceptable 3D structure was predicted, and the Ramachandran plot analysis has shown that most amino acids are in the correct positions (99 percent). There is no need to enhance or organize the analysis for further evaluation (figure 1).

**Figure 1.**
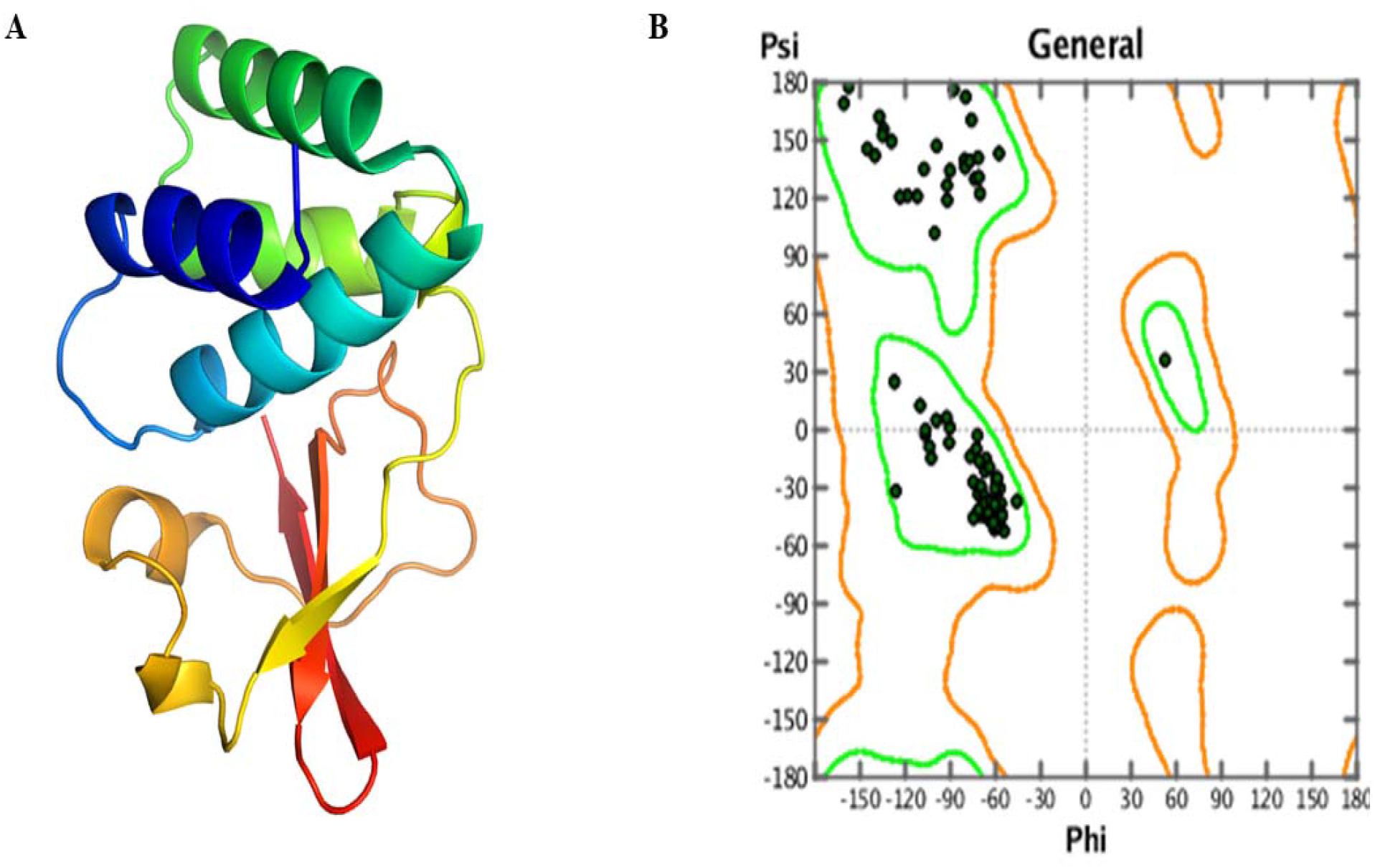
The graphical depiction of VP35 structure (A) shows the 3D structure MARV VP35 (B) A Ramachandran plot analysis reveals that most amino acids are located in favourable areas.

### Pharmacophoric based virtual screening from ZINC database

In CADD, virtual screening is a rapid and effective method to find novel therapeutic molecules. The ZINC pharmacophore programme had been used to filter the ZINC database’s millions of drug-like combinations. Hydrophobic (X= 3.97, Y= -0.14, Z= 0.00 and radius = 1.00), H acceptor (X= 2.1, Y= 0.25, Z= 0.00 and radius = 0.70), and H donor (X= 5, Y= -1.45, Z= 0.00 and radius = 1.00) characteristics were used to construct the pharmacophore complex model. The ZINC database yielded a high-quality inhibitor with characteristics comparable to the FGI-103 inhibitor (Figure 2). From the ZINC database, a total of 40570 chemicals were retrieved. After using Lipinski’s rule of five, only 2040 drug-like molecules were selected. The MOE program’s minimization technique was utilized to create a ready-to-dock catalogue.

**Figure 2.**
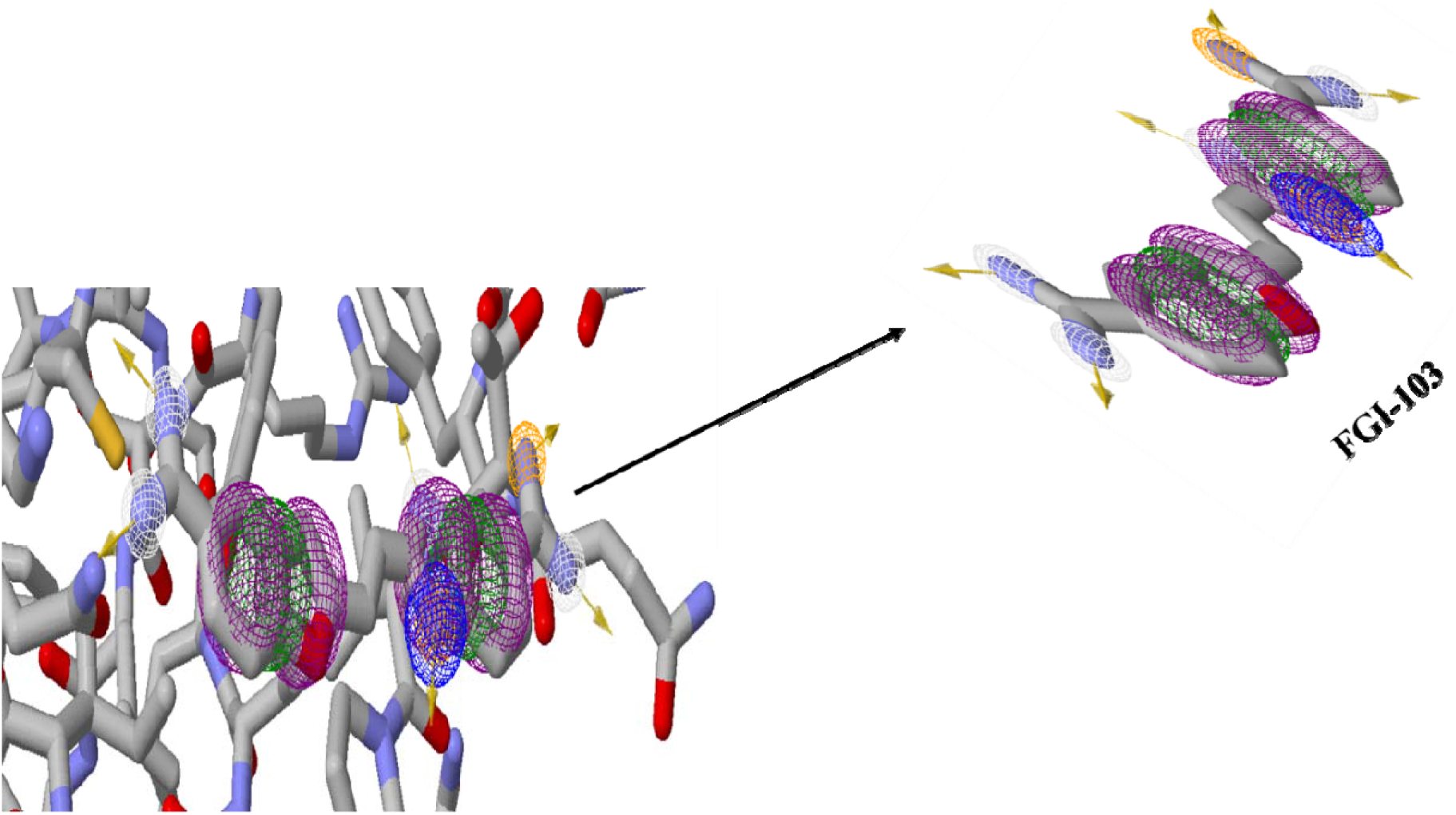
The ZINC pharmer programme produced a graphical depiction of the pharmacophoric combination (VP35 + FGI-103).

### Molecular docking

The significance of molecular docking has been determined in drug development. The MARV VP35 active site, the drug development portal, was found utilizing the active site discovery algorithm. MOE locked the VP35 protein, and the lower S-score and RMSD docking complex scored first. New drug-like compounds have been selected for screening with the lowest S-score (−11.23) and RMSD (2.4). The VP35 protein was also attached to a library of drug-like compounds. For further research, the top 250 small molecules with the lowest S-core and RMSD were selected. The binding relationship between these 250 chemicals and VP35 was determined using MOE Ligplot. To determine the bond strength of the bonds involved, the technique GB/VI Generalized Bond volume Required (MOE) was employed. Compounds with a higher interaction with MARV VP35 were chosen than the reference inhibitor, as indicated in Table 1. A total of 250 medications with 30 small molecules were tested for the ADMET profile.

**Table 1:**
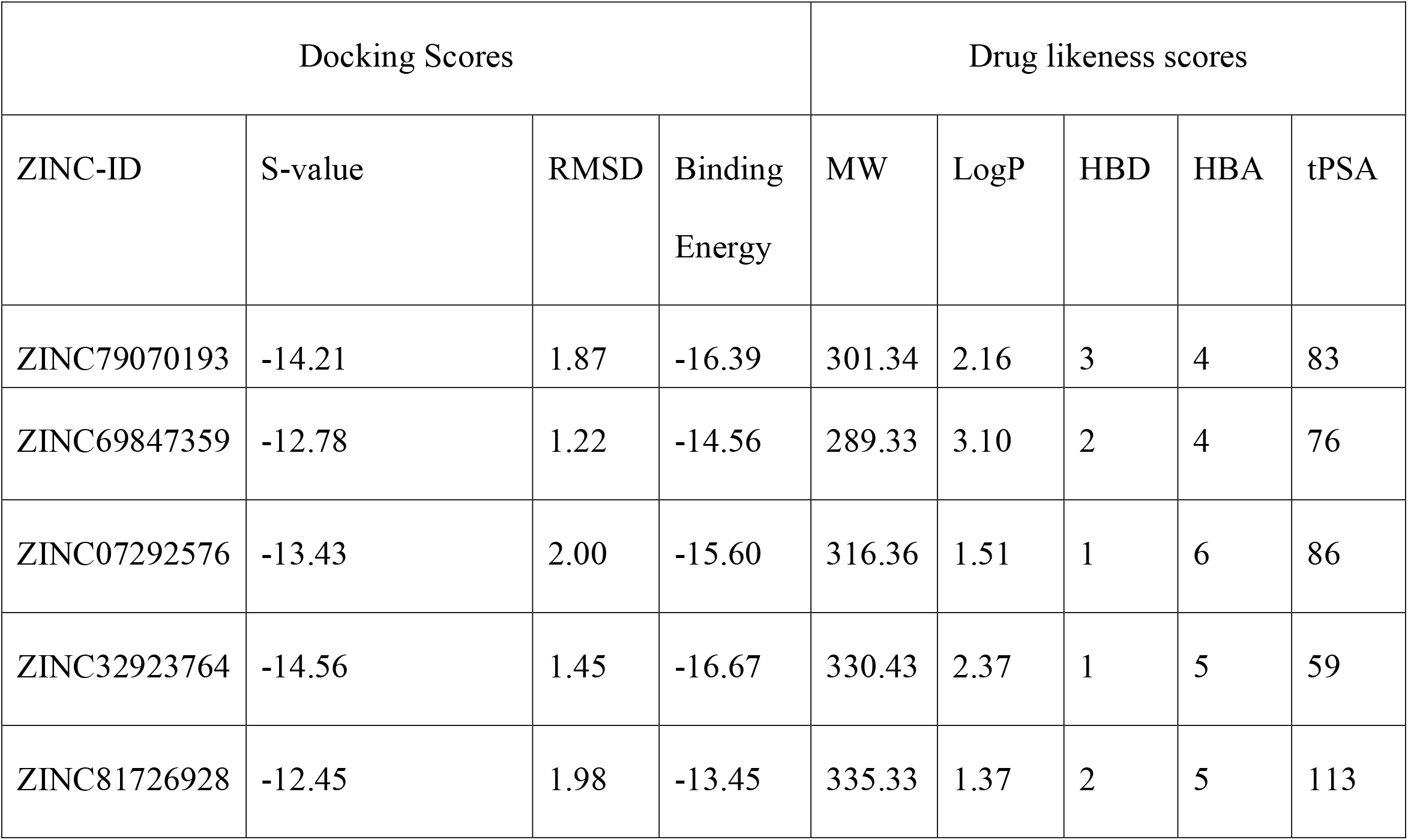
Properties of drug-like compounds and docking scores.

| Docking Scores |  |  |  | Drug likeness scores |  |  |  |  |
| --- | --- | --- | --- | --- | --- | --- | --- | --- |
| ZINC-ID | S-value | RMSD | Binding Energy | MW | LogP | HBD | HBA | tPSA |
| ZINC79070193 | -14.21 | 1.87 | -16.39 | 301.34 | 2.16 | 3 | 4 | 83 |
| ZINC69847359 | -12.78 | 1.22 | -14.56 | 289.33 | 3.10 | 2 | 4 | 76 |
| ZINC07292576 | -13.43 | 2.00 | -15.60 | 316.36 | 1.51 | 1 | 6 | 86 |
| ZINC32923764 | -14.56 | 1.45 | -16.67 | 330.43 | 2.37 | 1 | 5 | 59 |
| ZINC81726928 | -12.45 | 1.98 | -13.45 | 335.33 | 1.37 | 2 | 5 | 113 |

### Validation and ADMET analysis

The properties of ADMET on fifty drug-like compounds were investigated using the pkCSM server. Only five of thirty small drug-like compounds passed the ADMET test. A harsh gift for endothelial cells is the Blood-Brain Barrier (BBB), which stops the mind from getting medication. BBB plays an essential function in the creation of drugs (24,25). Oral bioavailability is an important consideration when choosing an active and successful medication for a particular patient (26,27). The characteristics of the ADMET analysis listed below are critical for finding successful therapeutic compounds; as a result, this parameter plays an essential role in the virtual screening of new medicines, such as compounds against a particular illness or pathogen.

The ADMET parameters are considered to include p-glycoprotein substrate/inhibitor, Blood-brain Barrier substrate/inhibitor, Human intestinal preparation, CaCo2 permeability, Renal organic cation transporter, CYPs enzyme, AMES toxicity. A good and effective medicine must satisfy the ADMET standard. Five compounds satisfied the Admet criteria out of fifty medicines that resembled them the most. These five drug-like small compounds (ZINC79070193, ZINC69847359, ZINC07292576, ZINC32923764, ZINC81726928) have substantially accepted the ADMET criteria (Table 2). For to choose therapeutic development molecules, the following criteria were used: ZINC ID, docking ranking, RMSD rating, binding energy value, VP35 binding residue, Lipinski’s findings (Table 1), and ADMET study ZINC ID, docking ranking, RMSD rating, binding energy value, VP35 binding residue, Lipinski’s results (Table 1), and ADMET study (Table 2). The chemicals chosen may be utilized as VP35 replication inhibitors that are new, structurally different, and possibly active (Figure 3).

**Table 2:**
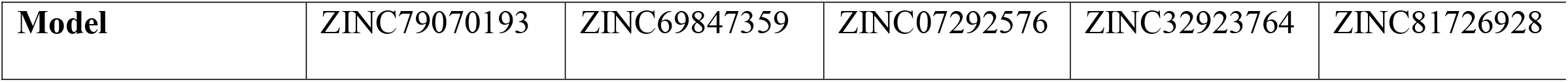
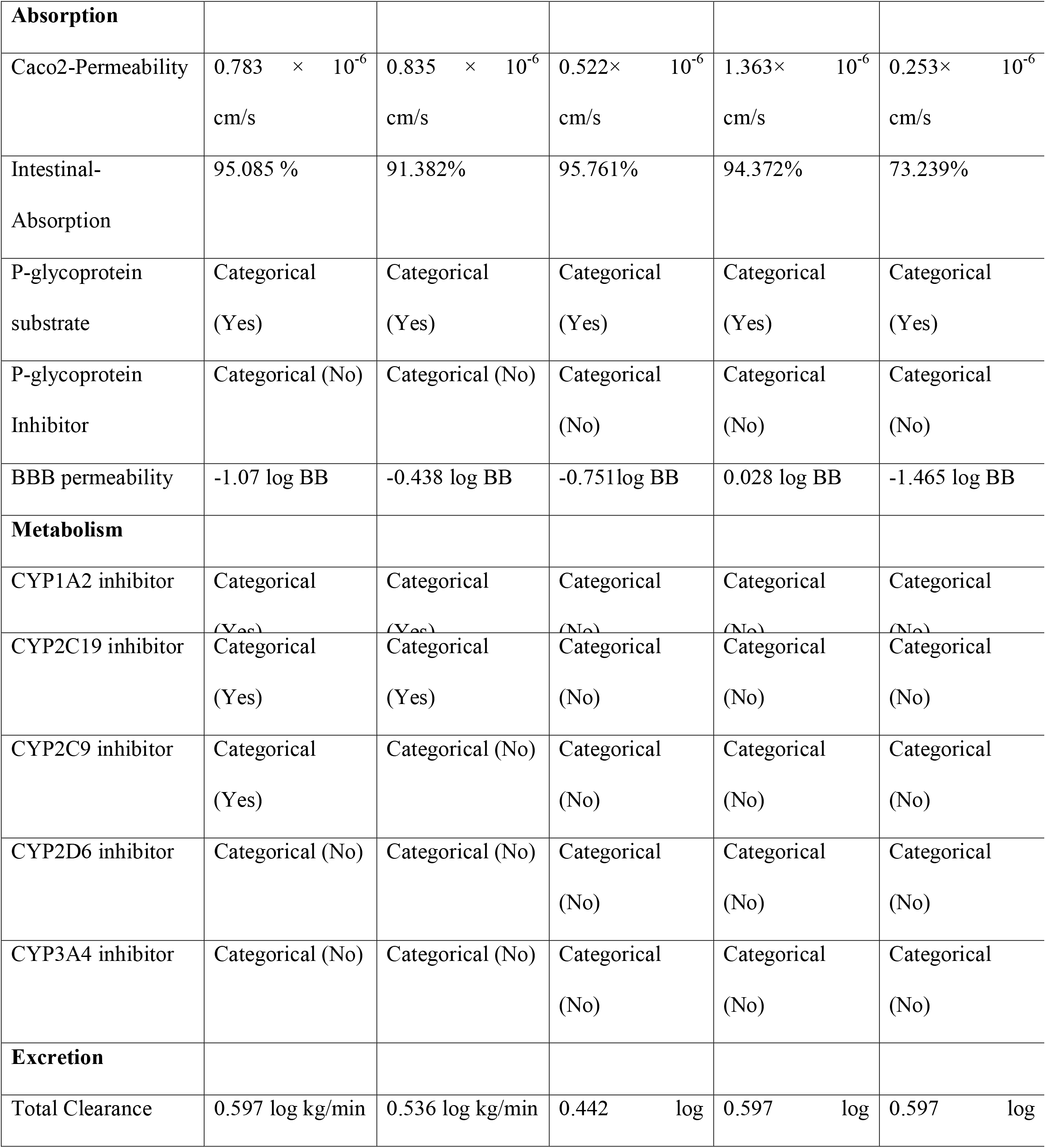
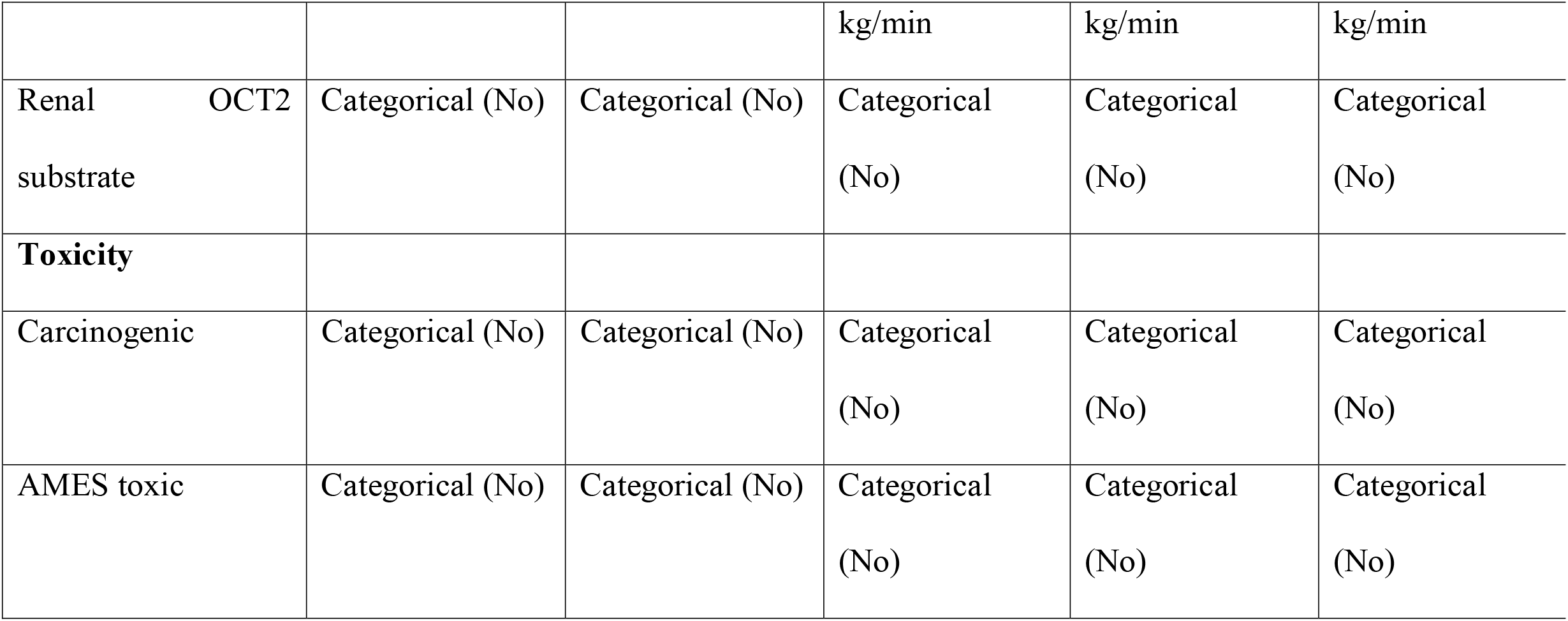
ADMET summary of drug compounds from ZINC database.

**Figure 3.**
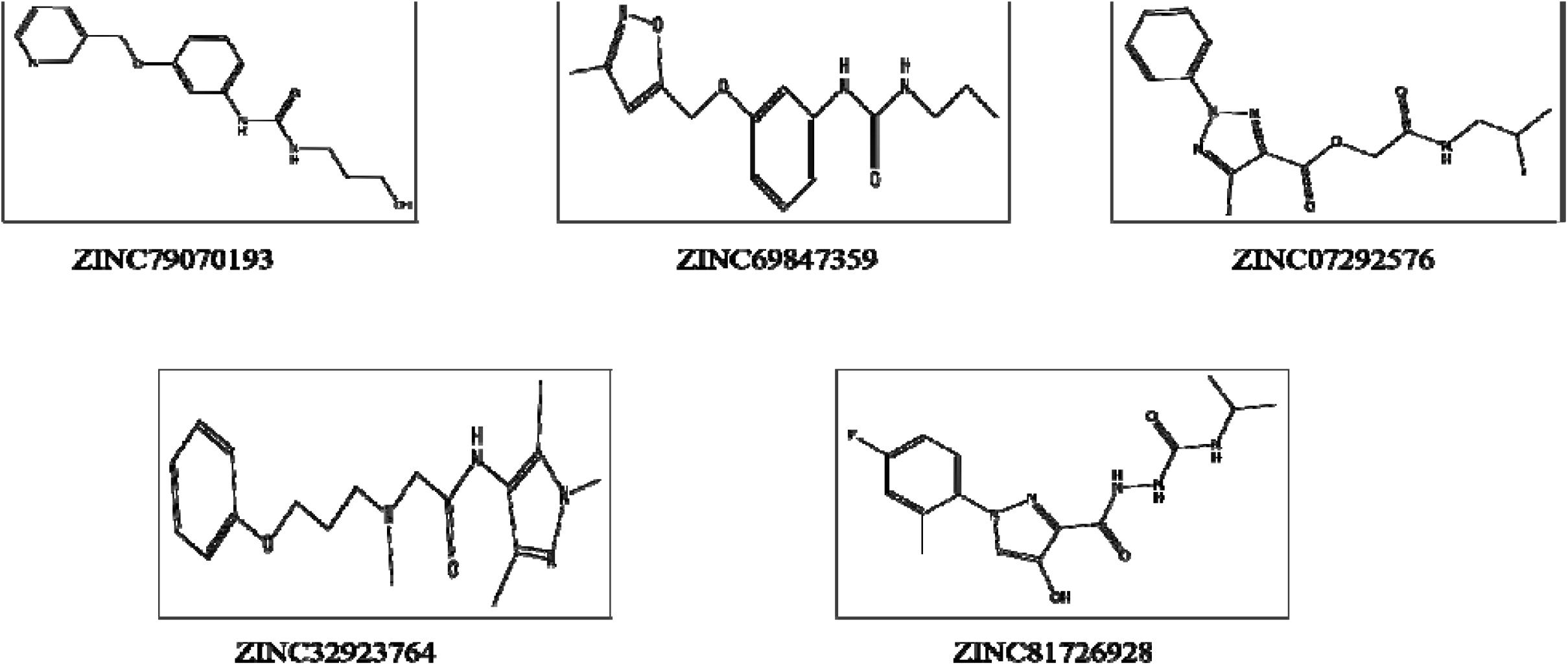
Representation of the new drug-like compounds’ 2D structures.

### Receptor-ligands interaction analysis

The intensity of the relationship between the amino acid receptor and the ligand atoms is measured by the S-docking score. Five small medications such as compounds were selected as novel effective molecules based on the score (S-value), the binding energy and other ADMET tests. Five medicinal compounds such as ZINC79070193, ZINC69847359, ZINC07292576, ZINC32923764, and ZINC81726928 produced a substantial complex of interactions with MARV VP35 active amino acids. Figure 4 shows the matching conformation of the ligand 3D docking and the proposed method of interaction with VP35. In total, A) LYS-237, 327, 328, 337, SER-271, 371 and ARG-271, which were involved in hydrogen bonding and polar interaction with Ligands, were the most active amino acids found.

**Figure 4.**
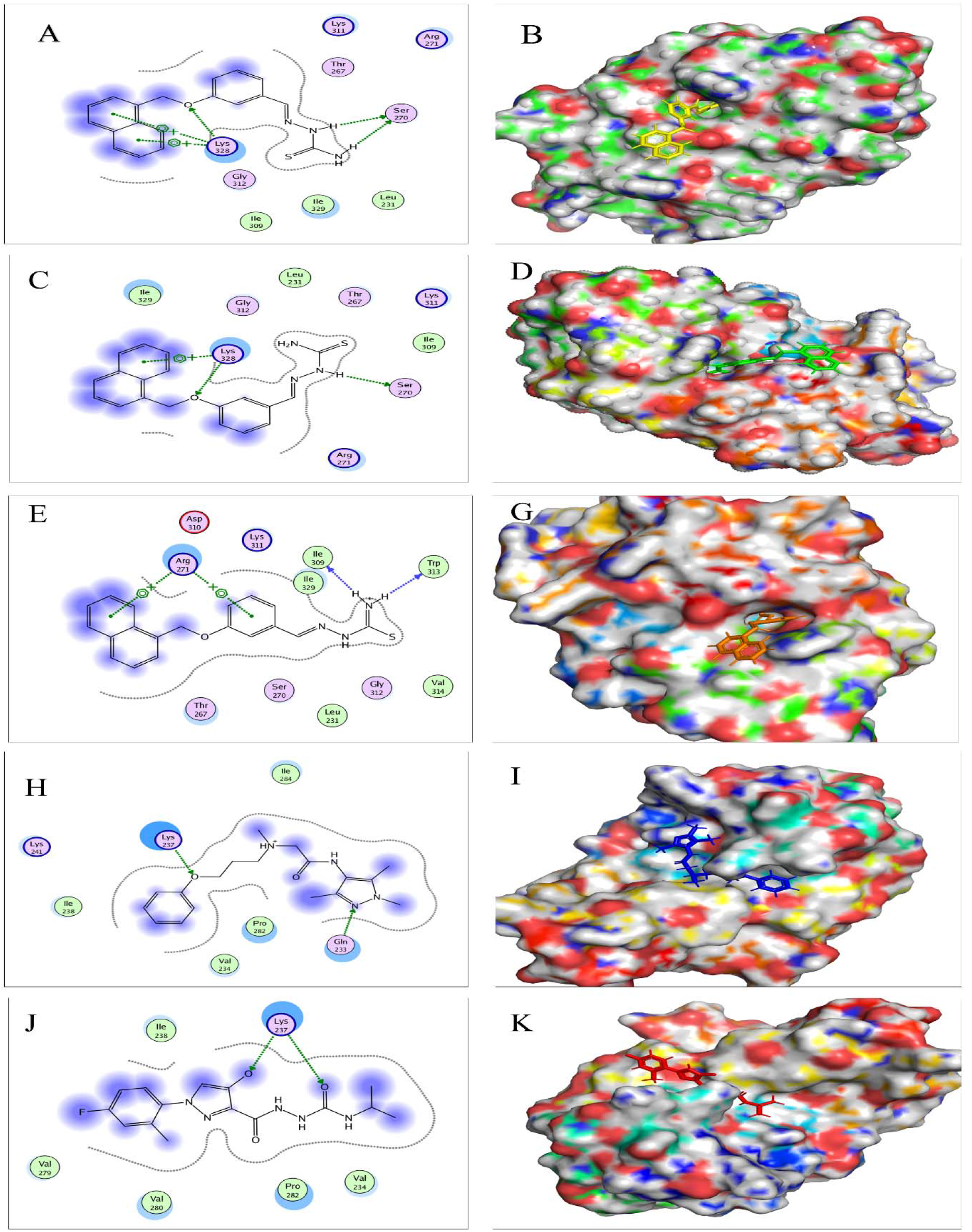
The top five drugs, such as compounds, were docked with the MARV target VP35 protein. (**A**) The complex’s 2D conformation shows that LYS-328 and SER-371 establish hydrogen bonds with the ligand ZINC79070193. The docking posture of ZINC79070193 (Yellow hue) to the VP35 protein is shown in (**B**). (**C**) The 2D conformation of the complex shows that LYS-328 and SER-271 establish hydrogen bonds with the ligand ZINC69847359, while ARG-271, LYS-311, THR-267, and GLY-312 are engaged in the interface accessible region (**D**). (**E**) The complex’s 2D conformation shows that ARG-271 forms a hydrogen bond with the ligand ZINC07292576, whereas ILE-309, 329, and TRP-313 interact polarly. The docking posture of ZINC07292576 (orange colour) to the VP35 protein is shown in (**F**). (**G**) The 2D conformation of the complex shows that LYS-337 and GLY-233 establish hydrogen bonds with the ligand ZINC32923764, while VAL-234, PRO-282, ILL-238, and LYS-341 are engaged in the interface accessible region (**H**). (**I**) The 2D conformation of the complex shows that LYS-237 forms a hydrogen bond with the ligand ZINC81726928 and that PRO-282, VAL-234,279. 280, and ILE-238 are engaged in the interface accessible region (**J**).

## CONCLUSION

In the quest for a MARV vaccine, many experimental trials have been undertaken. However, no medication or vaccine is presently being developed to control this disease effectively. As a consequence, developing a low-cost MARV-controlling antiviral drug is essential. Novel medication-like small compounds are effective against viral illnesses thanks to a gradual strategy for drug development.

Traditional drug development techniques are ineffective and time-consuming. Because of this, the primary objective of the present study was to use the ZINC database for pharmacopoeia simulation scanning, molecular docking of selected drugs, and binding testing of MARV VP35 interaction. The MARV VP35 active site was well connected with the five chemicals ZINC79070193, ZINC69847359, ZINC07292576, ZINC32923764, ZINC81726928. These results point to the possibility of these chemicals being utilized as a MARV medication. It would be beneficial in vivo drug research and development.

